# Comparative analysis of mitochondrial genome between UG93A and UG93B reveal common feature of 5’-end heterogeneity in mitochondrial genes of kenaf

**DOI:** 10.1101/523647

**Authors:** Xiaofang Liao, Yanhong Zhao, Aziz Khan, Xiangjun Kong, Bujin Zhou, Min Li, Meiling Wei, Shuangshuang Peng, Fazal Munsif, Ruiyang Zhou

## Abstract

Plant cytoplasmic male sterility (CMS) being maternal phenomenon trait that result from pollen abortion and closely linked with mitochondrial DNA rearrangement in many crops including kenaf. However, the molecular mechanism in kenaf is poorly known. In present work, we described the mitochondrial genome in isonuclear CMS line UG93A and its maintainer line UG93B. Findings of the current study revealed that a total of 398 SNPs and 230 InDels were identified in UG93A mtDNA. Total of 26 SNPs variations and three InDels were identified in the coding region of *atp6*, indicating its active role in mitochondrial genome re-arrangement. Northern blot analysis showed that the transcripts of *atp1, atp4, atp6, cox3* and *sdh4* in F_1_ were consistent with UG93A but different for UG93B. The transcript of *atp9* was found similar between UG93B and F_1_ while different for UG93A, which depict that *atp9* may be regulated by nuclear genes in F_1_ hybrid. The expression of *atp9* in UG93A was substantially lower compared with UG93B, suggesting its key role for energy supplying in microspore development of kenaf. Circularized RNA (CR)-RT-PCR revealed that mitochondrial RNAs with heterogeneous 5’-ends but uniform 3’ - ends are common feature in kenaf mitochondrial genes, and the promoter architecture analysis showed that the promoter sequences in kenaf mitochondrial genome are highly diverged in comparison to those in other plants. Our data highlight that the translation of mitochondrial genes in kenaf is closely associated with heterogeneity of the 5’-end of plant mRNA. The present result provides the basic information in understanding the transcription of kenaf mitochondrial genome and can be used as reference in other plants.

## Introduction

Angiosperm mitochondrial (mt) genomes are complex and differ substantially in size ranging from 208 kb in *Brassica hirta* (Kubo *et al.*, 2000) to 11.3 Mb in *Silene conica* (Sloan *et al.*, 2012). The mt genomes of angiosperms usually not only reveal phylogenetic relationships among plant species but also intraspecific differentiation in cytoplasm (Fujii *et al.*, 2010). The *Cucurbitaceae* genomes ranges from 380 kb (*Citrullus lanatus*) to 2,400 kb (*Cucumis melo*) due to the amount of chloroplast sequence (113 kb) transfers and short sequences duplications (370 kb) in larger *Cucurbita* mitochondrial genome (Alverson *et al.*, 2010; Rodriguez-Moreno *et al.*, 2011). In CMS mitochondrial genomes, LD-CMS, derived from *Oryza sativa*, has a size of 434 kb while that of CW-CMS has 559 kb derived from *Oryza rufipogon* due to genomic dynamic rearrangements (Fujii *et al.*, 2010). Despite substantial variation in size and physical mapping properties, plant mitochondria exhibited significant conservation in angiosperm genes, including 37-83 protein coding, tRNA and rRNA genes (Adams *et al.*, 2002). The shuffling of mitochondrial DNA(mtDNA) sequences by recombination, repeat sequences and noncoding sequences has vital in mt genome evolution by altering gene organization and creating chimeric genes (Hanson and Bentolila, 2004; Chen and Liu, 2014).

Cytoplasmic male sterility (CMS) is a common phenomenon causing incompatibility between nucleus and mitochondria and has been found in various plant species (Liu *et al.*, 2011; Luo *et al.*, 2013; Okazaki *et al.*, 2013; Heng *et al.*, 2014; Reddemann and Horn, 2018). CMS systems are not only useful genetic tool for hybrid crop breeding, but also ideal models for studying the genetic interaction and cooperative function of mitochondrial and nuclear genomes in plants (Hanson and Bentolila, 2004). Plant CMS may be caused by mutations, rearrangements or recombinations in the mitochondrial genome and novel chimeric genes deduced from mtDNA rearrangements have been identified in several plants (Hanson and Bentolila, 2004). In rice WA-CMS, a new mitochondrial chimeric protein WA352, interacts with the nuclear-encoded mitochondrial protein COX11, inhibiting its function in peroxide metabolism and triggering premature tapetal programmed cell death, leading to pollen abortion (Luo *et al.*, 2013). A CMS protein, ORFH79, binding to complex III, decreases its enzymatic activity in mitochondria and increases the reactive oxygen species (ROS) content through interactions with P61, resulting in pollen abortion in rice HL-CMS (Wang *et al.*, 2013). A toxic protein *ORF288*, co-transcribe with *atp6* and make transgenic plant fail to anther development, indicating its key role in *Brassica juncea* male sterility (Jing *et al.*, 2012, Heng *et al.*, 2018).

Mitochondria, chloroplast and other organelles contain its own genomes, however, their gene expression systems are distinct from other eukaryotes. Recently, studies have documented that the significance of regulating mitochondrial genes expression at RNA level, including regulation of multiple transcription initiation site at 5’-end of plant mitochondrial genes (Kuhn *et al.*, 2002; Kuhn *et al.*, 2005), formation of the 5’ - and 3’ - ends of mitochondrial mRNA, stabilization of stem-loop structures and degradation of polyadenylation of mitochondrial mRNA (Kuhn *et al.*, 2001; Forner *et al.*, 2007). Several nuclear-encoded proteins regulated plant mitochondrial genes expression. The pentatricopeptiderepeat (PPR) protein of *PPR336* gene has been shown to closely related with polysomes in mitochondria, which could be involved in translation of plant mitochondrial genes (Uyttewaal *et al.*, 2008). Another PPR protein of *MPPR6* gene that interacts with 5’-UTR of *rps3* mRNA is involved in 5’ maturation and translational initiation of *rps3* mRNA in maize. Functional deficiency of *MPPR6* results in a considerable reduction of *rps3* translation (Manavski *et al.*, 2012).

Kenaf (*Hibiscus cannabinus*) is an important fiber crop that is widely used in paper-making and weaving (Monti and Alexopoulou, 2013). Hybrid kenaf has attracted great interest due to its higher yield with better fiber quality and stress resistance (Tao *et al.*, 2008; Zhou *et al.*, 2008). The first cytoplasmic male sterile line in kenaf was bred from wild UG93, which then used as common way to produce F_1_ hybrid seeds (Zhou, 2002). CMS is an important factor in plant heterosis utilization and its mechanism of cytoplasmic male sterility is closely related to mt genome (Hanson and Bentolila, 2004). Sequencing of kenaf mt genome would not only be helpful for studying the evolutionary processes but could also clarify the inheritance mechanism for CMS in kenaf and the first mitochondrial genome of kenaf has been sequenced by Liao *et al.* (2018). Despite increasing interest in kenaf CMS, not much is known about its molecular mechanisms (Zhao *et al.*, 2013; Zhao *et al.*, 2015; Liao *et al.*, 2016; Zhao *et al.*, 2016). Efforts should be made to elucidate the factors underlying the mechanism of this important trait for hybrid breeding in kenaf. In present study, sequencing of UG93A mt genome in kenaf was determined and Northern blot analyses were used to compare differential transcription among UG93A, UG93B and F_1_ (UG93A/UG93R). Circularized RNA (CR)-RT-PCR was used to analyze the multiple transcription initiation sites of mitochondrial genes. Our data provides a better understanding of evolutionary processes of kenaf mt genome. Furthermore, this study could offer useful information in understanding the transcription of kenaf mitochondrial genome and can be used as reference in other plants.

## Materials and methods

### Plant materials

Kenaf CMS line UG93A is a naturally mutant of UG93 variety. It was provided by Institute of Bast Fiber Crops, Chinese Academy of Agricultural Sciences. The maintainer line UG93B was bred from wild-type variety UG93, and F_1_ (UG93A/UG93R) and was descendant of restorer UG93R hybridized with UG93A. These lines were belong to same nuclear background (UG93 nucleus) and hence considered the idealist materials for nuclear-cytoplasmic interaction investigations. This can assist in elimination of redundant genetic information from different nuclear background.

### Mitochondrial DNA isolation and Illumina Hiseq 2000 resequencing

Mitochondria were isolated from kenaf CMS line UG93A and purified from 7-day-old etiolated seedlings using differential centrifugation and discontinuous (18%, 23% and 40%) Percoll density gradient centrifugation according to Wilson and Chourey method (1984). Mitochondrial DNA isolation was performed according to Porebski *et al.* (1997) with modifications. Purified mitochondria were lysed with cetyltrimethylammonium bromide (CTAB) supplemented with 2% polyvinylpyrrolidone and 0.7% β-mercaptoethanol (Solarbio, Beijing, China) at 65 °C for 30 min. The lysis solution was extracted 2 to 3 times with chloroform/isoamyl alcohol (24:1), and absolute ethyl alcohol was used to precipitate mtDNA. DNase-free water (50 μL) was added to resuspend DNA pellets. The integrity, quality and concentration of UG93A mtDNA were analyzed using agarose gel electrophoresis, a NanoDrop 2000 (Thermo Scientific, Massachusetts, USA) and a Qubit fluorometer (Thermo Scientific, Massachusetts, USA), respectively. Extracted mtDNA of UG93A was prepared for next-generation sequencing (NGS) as follows: 5 μg of UG93A mtDNA was randomly sheared into fragments using a Covaris S220 (Thermo Scientific, Massachusetts, USA) with an insertion size of approximately 200-500 base pairs (bp) to construct a paired-end sequencing library. Fragmented DNA was combined with End Repair Mix (Qiagen, Duesseldorf, Germany), followed by the addition of A at the 3’ - end, and Illumina paired-end adaptor oligonucleotides were ligated to the sticky ends. The ligation mixture was purified with a QIAquick PCR Purification Kit (Qiagen, Duesseldorf, Germany). Several rounds of PCR amplification with the PCR Primer Cocktail and PCR Master Mix (Qiagen, Duesseldorf, Germany) were performed to enrich the adapter-ligated DNA fragments. DNA Clusters of PCR colonies were resequenced on a HiSeq 2000 sequencing platform using the recommended protocols from the manufacturer.

### Genome annotations and analyses

Low-quality sequences in raw data were filtered with the FASTX toolkit (http://hannonlab.cshl.edu/fastx_toolkit), and clean reads were obtained according to Cox *et al* (2010). BLASTn analysis was performed to exclude contamination of the chloroplast and nuclear genome sequences. Sequences were assembled into contigs using Velvet and Oases software (Cox *et al.*, 2010). Clean reads were mapped and aligned onto the reference mtDNA sequence of UG93B with BWA software (Li and Durbin, 2009). Genes encoding mitochondrial proteins and rRNAs were annotated using BLASTn and MITOFY (Alverson *et al.*, 2010) based on the known annotation of kenaf mitochondrial genes of UG93B. The tRNA genes were identified using tRNA scan-SE software (http://lowelab.ucsc.edu/tRNAscan-SE/).

### RNA Extraction and Northern Blot analysis

Total RNAs were isolated from anthers at binucleate stage using a Quick RNA Isolation Kit (Hua yue yang, Beijing, China) according to the protocol supplied by the manufacturer. For RNA blotting analysis, approximately 30 μg of total RNA was denatured and separated on a 1% denaturing formaldehyde agarose gel and transferred to a Hybond-N^+^ nylon membranes (GE Healthcare Life Sciences, London, UK). PCR products were labeled with DIG-High Prime DNA Labeling and Detection Starter Kit II (Roche, Mannheim, Germany), and the operating procedures according to the protocol supplied by the manufacturer of DIG-High Prime DNA Labeling and Detection Starter Kit II (Roche, Mannheim, Germany). The prober primes are shown in supplementary (Table S3).

### Synthesis of cDNA and qRT-PCR analysis

Two developmental stages of buds (at uninucleate stage and binucleate stage) were chosen to analyze the expression of *atp1, atp4, atp6, atp9* and *cox*3. Total RNA was isolated from the anthers of UG93A, UG93B, and F_1_ (UG93A/UG93R) with CTAB as described by Liu and He (2006). One microgram of total RNA was used in the reverse transcription reaction using the TransScript One-Step gDNA Removal and cDNA Synthesis SuperMix (TransGene Biotech, Beijing) to obtain 20 μL of the resulting cDNA solution. The gene-specific primers for qRT-PCR were designed by Primer Premier 5.0 software. Quantification was performed with SYBR^®^ Premix Ex Taq™ (Tli RNaseH Plus) (Takara, Beijing, China) using a Bio-Rad CFX96 instrument (Bio-Rad, California, USA). The housekeeping gene *Histone H3* (*Histone3*) of kenaf served as the internal reference. PCR cycling was denatured using a program of 95°C for 10 s and 40 cycles of 95°C for 5 s and 60°C for 30 s. For each sample three biological and technical replicates were used. The relative expression was calculated according to formula 2^−ΔΔCt^ for UG93A, UG93B, and F_1_ (UG93A/UG93R), with maintainer line UG93B as control. Primes for qRT-PCR were shown in supplementary Table S4.

### Statistical analysis

Data on relative genes expression of both CMS and maintainer line as well as F_1_ hybrid were statistically analyzed according to the procedure appropriate for one way analysis of variance using SPSS17.0 software.

### CR-RT-PCR analysis

The processing of RT-PCR analysis of circularized RNA was described by Kuhn *et al.* (2002). In brief, 1 μg of total RNA was self-ligated with T4 RNA ligase in a total volume of 10 μL under the conditions recommended by the enzyme manufacturer (Thermo Fisher Scientific, USA). First-strand synthesis was performed with primer AP1 and superscript reverse transcriptase on 1 μL ligated RNA following the instructions of the respective manual (TransGene Biotech, Beijing, China). DNA fragments comprising the 5’-3’ ligated mRNA end were subsequently amplified with primer pairs SP1/AP2, followed by a second PCR with nested primers SP2/AP3 using amplification products of first PCR as a template (Fig. S1). The products were finally cloned into a linearized pEASY-T1 Cloning vector according to the standard procedures followed by sequence analysis of more than 15 cDNA clones. The location of the primers was given with respect to the translation initiation codon (+1). All numbering are in respect to the translation start codon (+1). Primes for CR-RT-PCR analysis are shown in supplementary (Table S5).

### Sequence analysis

The 5’ and 3’ termini of the transcripts were determined by comparison between the clone sequences and the target gene sequences using the DNAMAN software. The promoter architecture was described by Crooks *et al* (2004) and analyzed using the online software WebLogo (http://weblogo.berkeley.edu/logo.cgi).

## Results

### Resequencing analysis of mitochondrial genome in UG93A

mtDNA of UG93A was randomly sheared into fragments with an average length of 300 bp to construct a paired-end sequencing library and resequenced on a HiSeq 2000 sequencing platform. Clean data were obtained after filtering low-quantity, short reads and adaptors from raw data. The number of reads, read length and total bases were counted before and after filtration. A total of 3.7×10^5^ paired-end reads with a filtered ratio of 2.16% were obtained, as were 1×10^8^ total bases with a filtered ratio of 11.36%. Clean data were aligned to the reference mtDNA of UG93B using BWA software. The coverage of UG93A mtDNA was 99.2%, aligning to UG93B mtDNA, with a coverage depth of 179× (Table S1).

### Analysis and annotation of SNPs in UG93A mtDNA

A total of 398 SNPs in UG93A mtDNA with a reference mitochondrial genome sequence of UG93B were observed. The distribution rate and frequency of SNPs in the whole UG93A mtDNA were counted per unit of 50 kbp. Results showed that the frequency of UG93A mtDNA was diverse in various regions. The variable frequency of SNPs from 350-400 kb was the highest (1.94 bp/kb) with the main mutant genes of *rrnL* and *rrnS*. Most of SNPs variation occurred in non-coding regions, intergenic regions and introns, including 362 SNPs (accounted 90% of total SNPs) distributed intergenic and introns. While only 37 SNPs (accounted 10% of the total SNPs) distributed in the coding region of mitochondrial genes (Table S2). The amino acid annotation in the variation of the coding region was analyzed. These results showed five SNPs with synonymous mutations and thirty-two non-synonymous mutations in the gene coding region, resulting in changes of the amino residues encoded by eight mitochondrial genes. Twenty-six of these SNPs variations were found in the coding region of *atp6*, while only one or two SNPs among other genes were found, indicating that the coding region of *atp6* in UG93A was an active recombination region (Fig. 1, Table 2).

**Table 2.** Annotation of SNPs in protein-coding genes of UG93A mtDNA

| No. | Variable sites | RefSeq | Base mutation | Gene name | Amino acids | Mutated amino acids |
| --- | --- | --- | --- | --- | --- | --- |
| **1** | **92263** | **A** | **C** | ***rpl10*** | **I** | **L** |
| **2** | **134785** | **C** | **T** | ***sdh4<sup>a</sup>*** | **G** | **S** |
| 3 | 192259 | A | G | *cox1* | G | G |
| 4 | 193141 | C | T | *cox1* | F | F |
| 5 | 193750 | T | G | *cox1* | L | L |
| 6 | 193303 | C | A | *cox1* | I | I |
| **7** | **271723** | **T** | **G** | ***rps14<sup>a</sup>*** | **L** | **F** |
| **8** | **470838** | **C** | **G** | ***atp6<sup>a</sup>*** | **E** | **D** |
| **9** | **470847** | **C** | **A** | ***atp6*** | **E** | **D** |
| **10** | **470863** | **C** | **T** | ***atp6*** | **S** | **N** |
| **11** | **470885** | **A** | **G** | ***atp6*** | **Y** | **H** |
| **12** | **470896** | **T** | **C** | ***atp6*** | **K** | **R** |
| **13** | **470911** | **T** | **A** | ***atp6*** | **Q** | **L** |
| **14** | **470927** | **G** | **C** | ***atp6*** | **P** | **A** |
| **15** | **470935** | **C** | **T** | ***atp6*** | **G** | **E** |
| **16** | **471116** | **T** | **C** | ***atp6*** | **N** | **D** |
| **17** | **471119** | **A** | **G** | ***atp6*** | **F** | **L** |
| **18** | **471128** | **A** | **G** | ***atp6*** | **Y** | **H** |
| **19** | **471131** | **A** | **C** | ***atp6*** | **Y** | **D** |
| **20** | **471136** | **C** | **A** | ***atp6*** | **C** | **F** |
| **21** | **471148** | **A** | **G** | ***atp6*** | **L** | **T** |
| **22** | **471149** | **G** | **T** | ***atp6*** | **L** | **T** |
| **23** | **471166** | **T** | **G** | ***atp6*** | **E** | **S** |
| **24** | **471167** | **C** | **A** | ***atp6*** | **E** | **S** |
| **25** | **471177** | **T** | **A** | ***atp6*** | **L** | **F** |
| **26** | **4711182** | **G** | **A** | ***atp6*** | **H** | **Y** |

| Continue |  |  |  |  |  |  |
| --- | --- | --- | --- | --- | --- | --- |
| **27** | **471228** | **A** | **C** | ***atp6*** | **C** | **W** |
| **28** | **471232** | **C** | **A** | ***atp6*** | **G** | **V** |
| **29** | **471237** | **G** | **T** | ***atp6*** | **A** | **E** |
| **30** | **471261** | **A** | **T** | ***atp6*** | **F** | **L** |
| **31** | **471238** | **G** | **T** | ***atp6*** | **A** | **E** |
| **32** | **471273** | **G** | **A** | ***atp6*** | **S** | **F** |
| **33** | **471277** | **G** | **A** | ***atp6*** | **T** | **M** |
| **34** | **512025** | **A** | **C** | ***cox2*** | **T** | **T** |
| **35** | **512031** | **G** | **T** | ***cox2*** | **M** | **I** |
| **36** | **539731** | **C** | **G** | ***atp4*** | **I** | **M** |
| **37** | **568501** | **A** | **T** | ***rps7<sup>a</sup>*** | **F** | **L** |

Boldface represent nonsynonymous mutation. a, indicated the transcription of this gene occurs on the antisense strand.

**Fig. 1.**
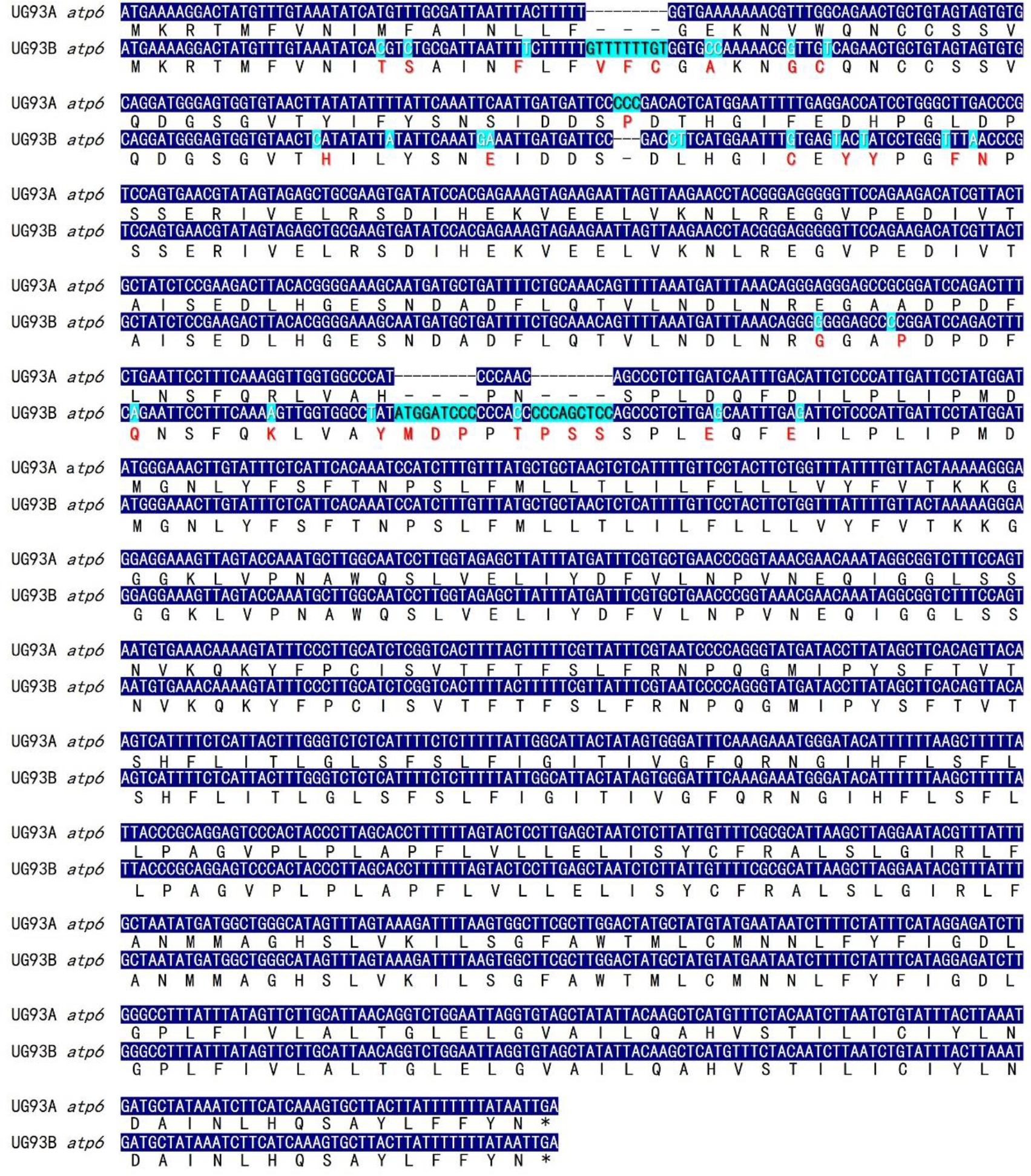
Alignment of the *atp6* between UG93A and UG93B. The variable nucleotide between UG93A and UG93B was marked with light blue background, and the corresponding variation of amino acid residues was indicated with red letter.

### Analysis and annotation of InDels in UG93A mtDNA

A total of 230 small fragments of insertion/deletion (InDel) variations were found in UG93A mtDNA compared to UG93B mtDNA, ranging from 1 bp to 31 bp (Fig. 2). The short InDels were mainly distributed from 1 bp to 5 bp in total of 178, accounting for 77.4% of total InDels, while 52 InDels were distributed from 6 bp to 31 bp, accounted for 22.6%. In particular, a total of 99 InDels with a size of 1 bp and 38 InDels with a size of 4 bp were distributed in UG93A mtDNA, accounting for 43% and 16% of the total InDels, respectively. Other InDels were rarely distributed from 2% to 6%. Most InDels were distributed between the intergene region and intron. Only 11 InDels were distributed in the coding region resulting variation in amino acids encoded by mitochondrial genes (Table 1).

**Fig. 2.**
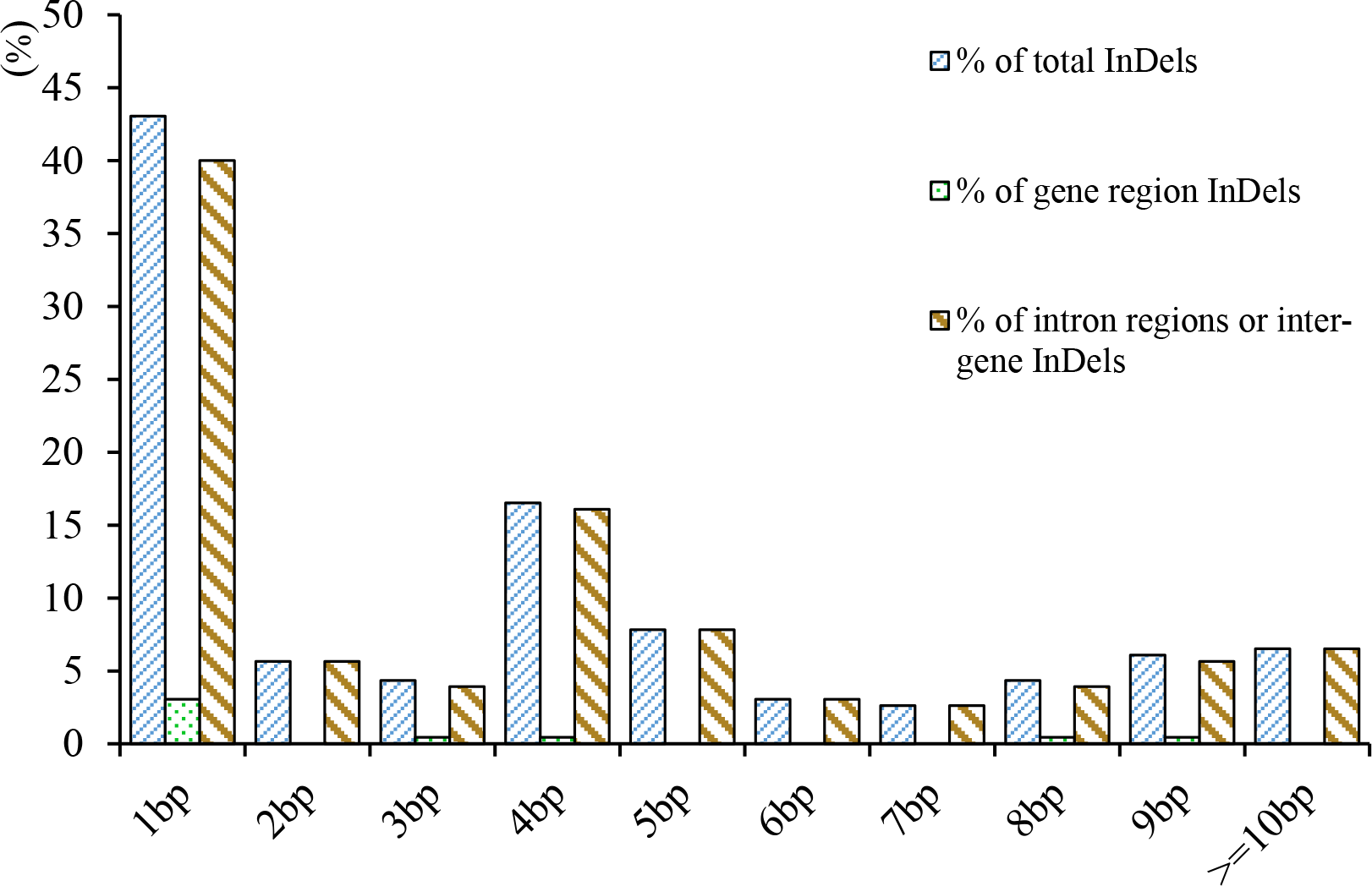
Distribution of different sized InDels in UG93A mtDNA

### Transcript analysis of kenaf mtDNA

To find the mitochondrial genomic regions that possibly contain the candidate genes for kenaf CMS, a total of 36 protein coding mitochondrial genes were used to design probe primers to examine the possible alteration in the mitochondrial transcription patterns among UG93A, UG93B and F_1_ (UG93A/UG93R) by Northern blotting analysis. Most of mitochondrial genes shared the same mRNA among UG93A, UG93B and F_1_ (UG93A/UG93R), excluding six genes, *atp1, atp4, atp6, atp9*, *cox3* and *sdh4* (Fig. 3). Moreover, these mitochondrial genes of *atp1, atp4, atp6, cox3* and *sdh4* showed the same transcription patterns between UG93A and F_1_ (UG93A/UG93R), but different from UG93B, indicating that the transcript patterns of these five mitochondrial genes were not regulated by nuclear genes in kenaf (Fig. 3). Furthermore, four probes of *atp1, atp6, cox3* and *sdh4* detected the mRNA patterns with strong signal among UG93A, UG93B and F_1_ (UG93A/UG93R), but weaker signal in UG93B with *atp4* as prob (Fig. 3b). The probe of *atp9* detected a special mRNA with relatively smaller size and weaker signal in UG93A and F_1_ (UG93A/UG93R) (Fig. 3d), This suggests that the differential transcript of *atp9* in F_1_ (UG93A/UG93R) may be regulated by nuclear genes despite the same cytoplasm between UG93A and F_1_ (UG93A/UG93R) and correlate to kenaf CMS.

**Fig. 3.**
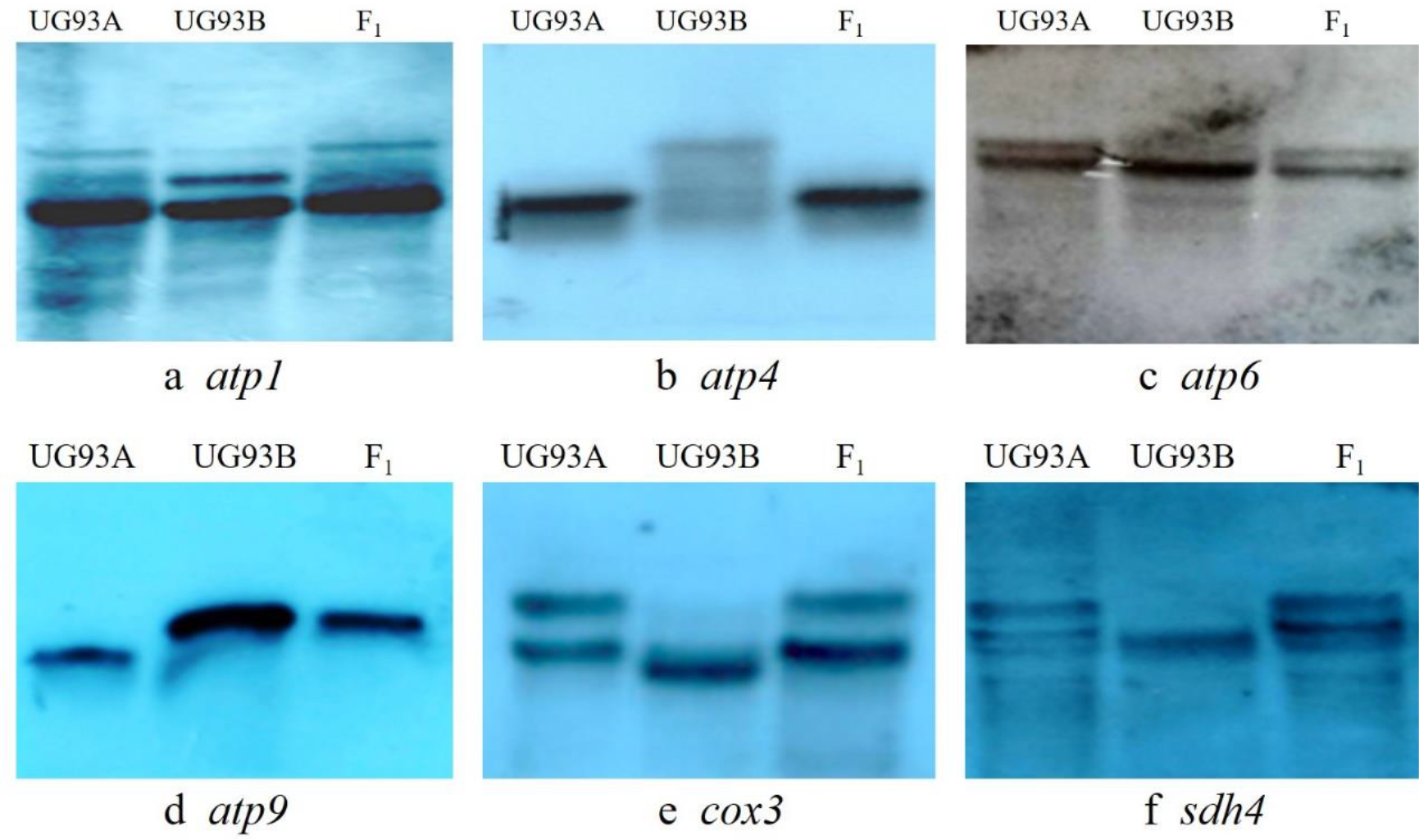
Transcriptional analysis of the kenaf mitochondrial genes.

### Relative expression of mitochondrial respiratory chain complex genes

Real-time quantitative PCR was performed for determination of relative expression levels of *atp1, atp4, atp6, atp9* and *cox3* in UG93A, UG93B and F_1_ (UG93A/UG93R) in the monocaryotic phase and dikaryophase of kenaf anther. The levels of *atp1, atp4* and *atp9* in the CMS line UG93A were lower compared with UG93B in both monocaryotic phase and dikaryophase. The expression of *atp9* in UG93A was significantly lower than in UG93B, indicating that *atp9* played a central role in the energy supply for the microspore development in kenaf (Fig. 4a), and the relative expression in F_1_ (UG93A/UG93R) was higher compared with UG93B at monocaryotic phase, while significantly lower at dikaryophase of anther development (Fig. 4a). The expression of *atp6* and *cox3* was significantly lower than UG93B and F_1_ (UG93A/UG93R) in the monocaryotic phase, but substantially higher than UG93B in dikaryophase (Fig. 4b). These data revealed that mitochondrial gene expression in kenaf may be regulated by multiple factors.

**Fig. 4.**
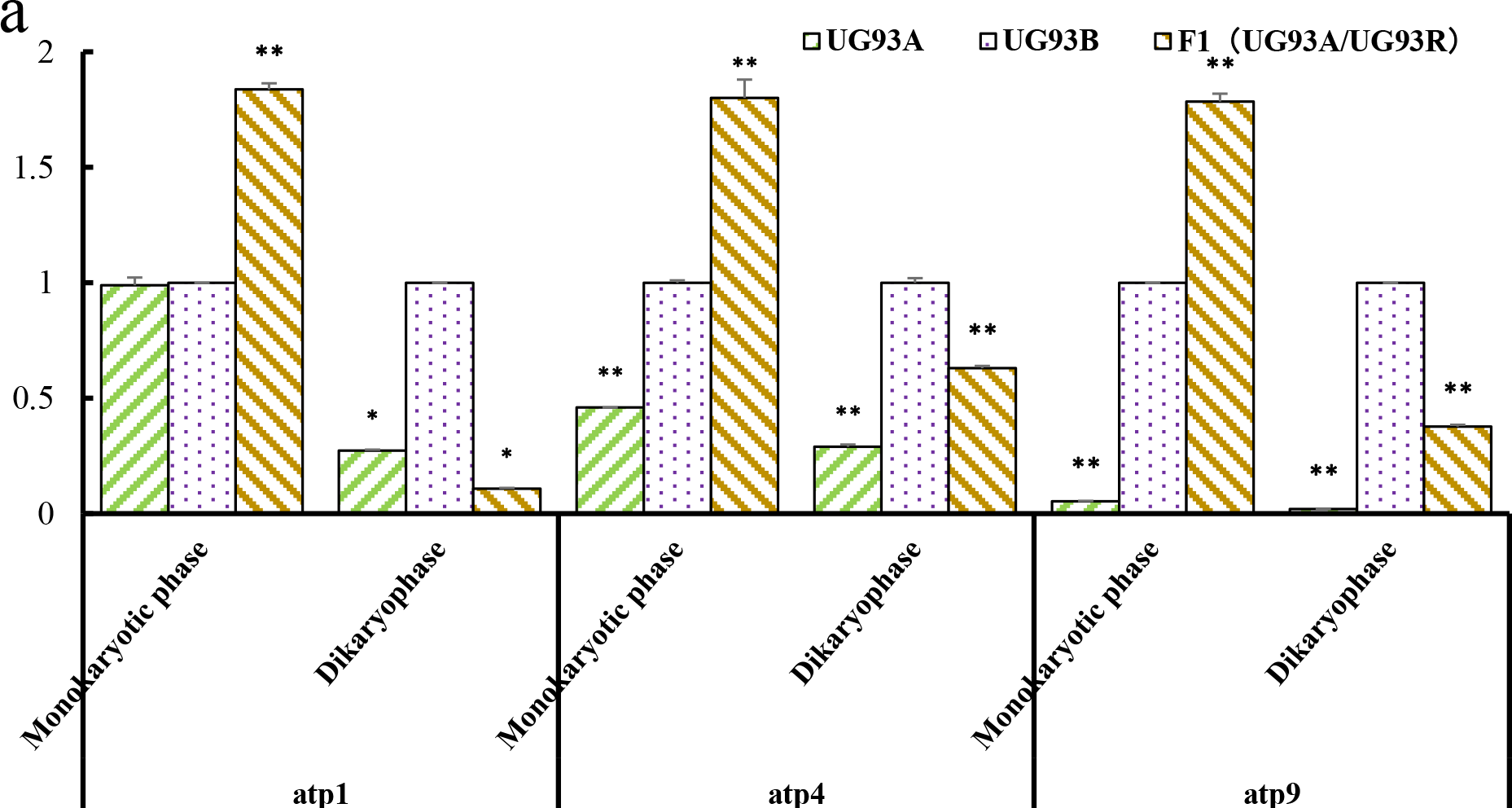
Relative expression analysis of *atp1, atp4, atp6, atp9* and *cox3*. The housekeeping gene *Histone3* was used as an internal control. Error bars represent standard deviation (n=3). The significance of differences was assessed by Student’s t-test (*P<0.05, **P<0.01).

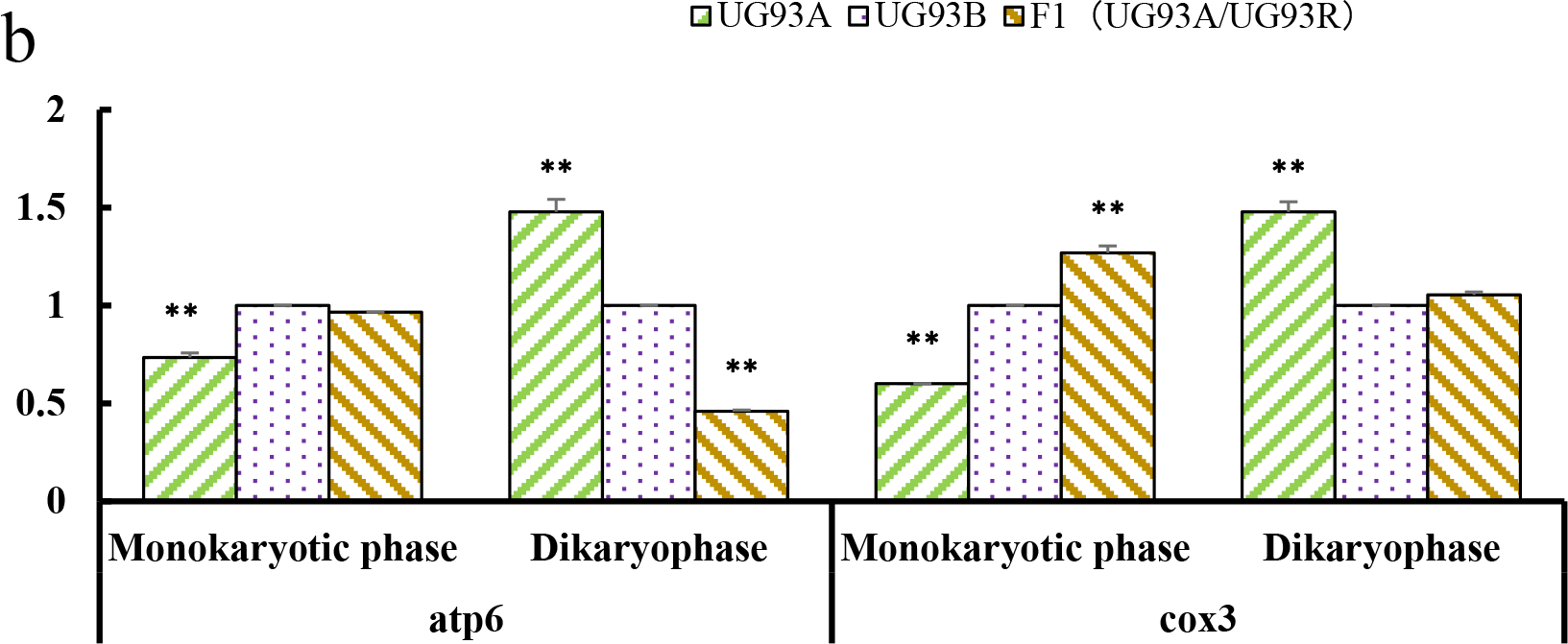

### Circularized RNA (CR)-RT-PCR analysis of mitochondrial genes

Previous analysis revealed that the transcripts of *atp1, atp4, atp6* and *cox3* can be transcribed into polycistronicm transcripts, but *atp9* with monocistronic transcripts. The transcripts of *atp1, atp4, atp6, atp9* and *cox3* were analyzed by CR-RT-PCR in UG93A, UG93B and F_1_ (UG93A/UG93R) (Fig. 5). Results showed multiple transcription initiation sites at the 5’ - ends upstream of *atp1, atp4, atp6* and c*ox3*, but the transcript termination sites of 3’ - ends were uniform in UG93A, UG93B and F_1_ (UG93A/UG93R) except for *atp9*. Sequences analysis revealed that the 3’-ends of *atp1* were uniform (+49 relative to the stop codon) in UG93A, UG93B and F_1_ (UG93A/UG93R). Three 5’- ends were located at −78, −138 and −213 relative to the start codon in UG93A. Only one 5’-ends was located at −176 in UG93B and two 5’-ends were located at −158 and −176 in F_1_ (UG93A/UG93R). Interestingly, the CR-RT-PCR profile was different slightly with the transcript of *atp1*. Two major transcripts were identified in UG93B, whereas only one 5’- end was identified by CR-RT-PCR, suggesting that 5’- end could be involved in the translational efficiency of plant mitochondrial RNA. 2-bp insertion at the (−11) − (−12) locus and nine nucleotide variations at the position (−92) − (−114) upstream of the start codon in UG93A and F_1_ (UG93A/UG93R), respectively (Fig. 6, Fig S2).

**Fig. 5.**
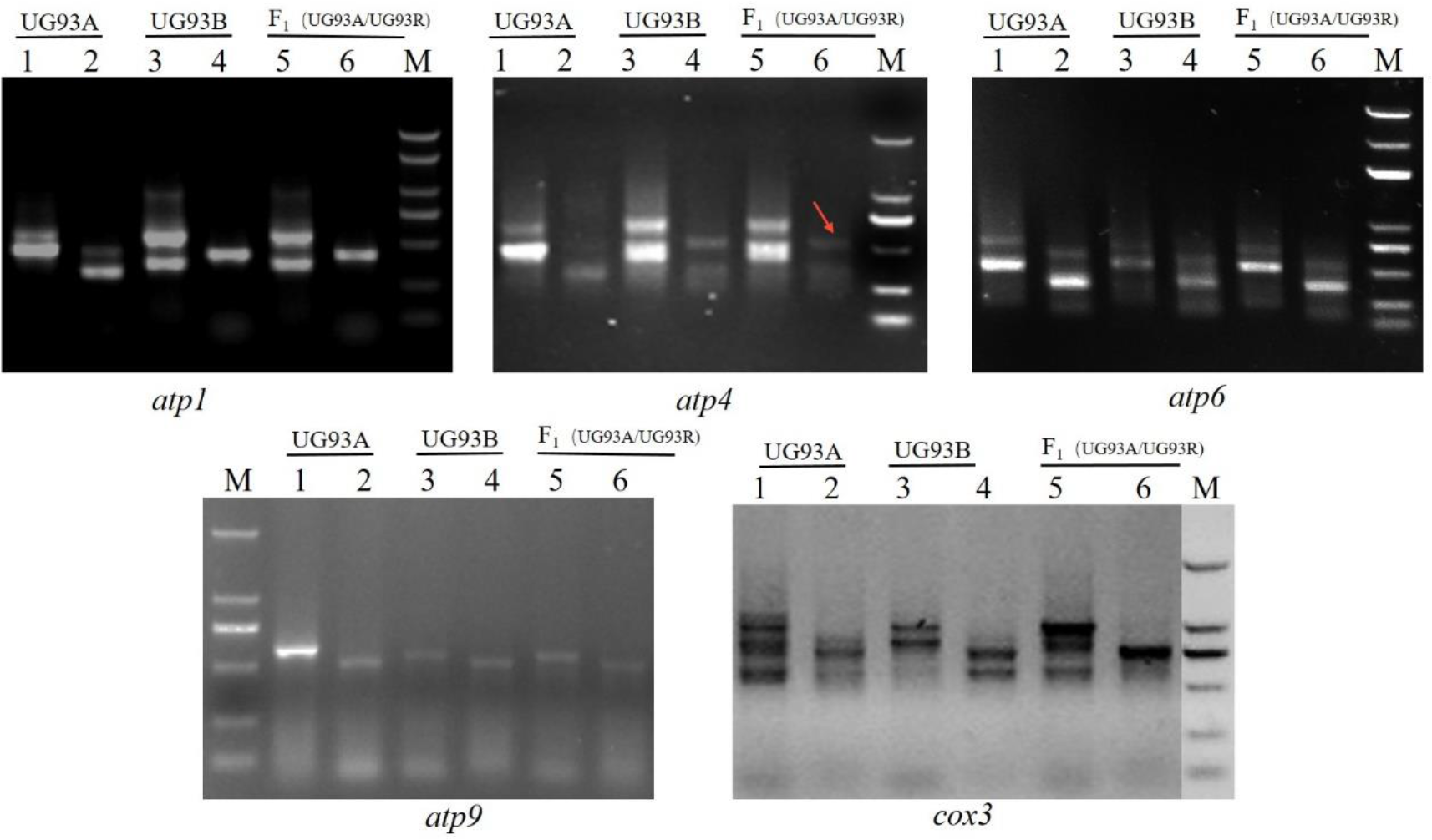
CR-RT-PCR analysis of mitochondrial genes. The second amplification product of *atp4* in F_1_ was given in red arrow.

Similar observations have been made in other mRNAs. There was only one 5’- end of *atp4* located at nucleotide position −307 upstream of the start codon in UG93A. Three 5’- ends located at nucleotide positions −56, −274 and −376 upstream of the start codon in UG93B and two 5’ - ends located at nucleotide position −74 and −307 in F_1_ (UG93A/UG93R). The 3’ - ends were uniform (+ 71 relative to the stop codon) in the three cytoplasm of kenaf (Fig. S3, S4).

**Fig. 6.**
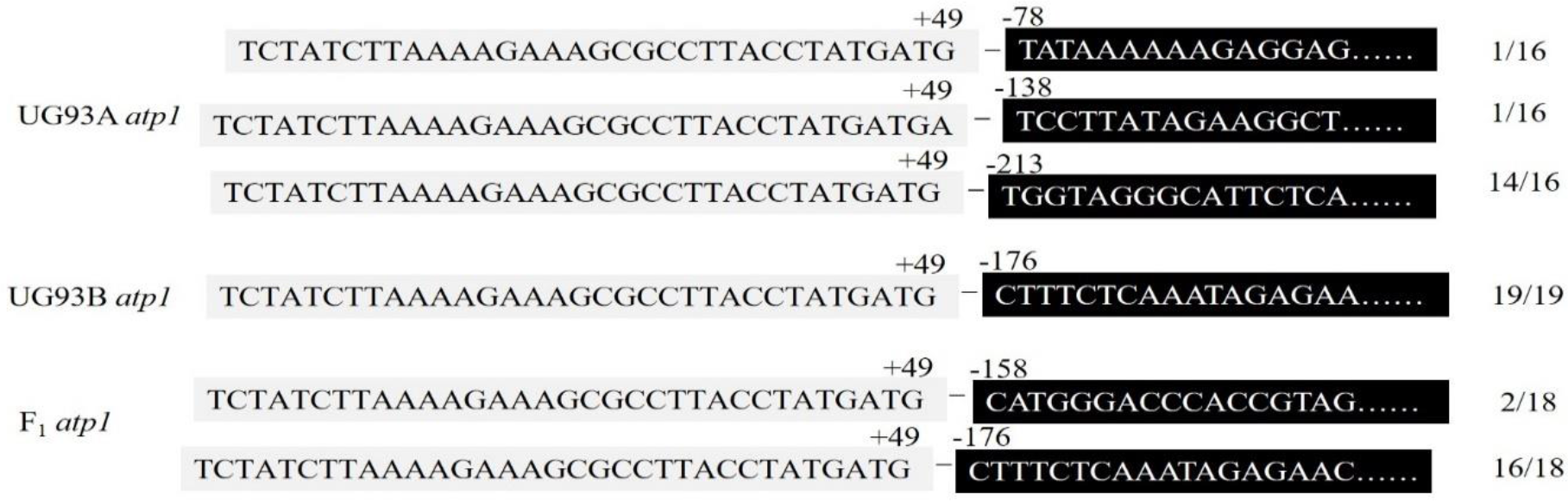
5’- and 3’- ends detected of *atp1* by the CR-RT-PCR. Sequences of the cDNA covering the 5’-3’ ligation site of *atp1* transcripts. The numbering of the 5’- end refers to the translation start codon and the 3’-numbers are relative to the stop codon. The numbers on the right indicate the frequency with which the distinct 5’ - and 3’ - ends are identified in individual clones. The ligation site is indicated by a horizontal dash. Sequences corresponding to the 5’-terminal part are given in white letters in black boxes.

The transcript characteristics of *atp6* in F_1_ (UG93A/UG93R) was identity with UG93A, located at nucleotide positions −130 and −381 upstream of the star codon, respectively, and the transcription initiation site in UG93B was located at nucleotide positions −130 and −310 upstream of the start codon, respectively. The 3’ - ends were mapped at the same position in the three cytoplasm of kenaf (+ 42 relative to the stop codon) (Fig. S5). Compared with UG93B, one-bp insertion and a nine-bp deletion was identified at nucleotide position −26 and (−49)−(−57) upstream of the reading frame, respectively. Nucleotide variation happened from the position −229 upstream of the reading frame in UG93A, UG93B and F_1_ (UG93A/UG93R). These results suggested that they were the active region of mitochondrial genome recombination (Fig. 7, Fig. S6). In the cDNA sequence of *atp6*, an RNA editing with C-T conversion occurred at the nucleotide position +1183 of reading frame in UG93A or +1204 of reading frame in UG93B, resulting in generation of a stop codon TAA, that lead to early termination of the CDS region of *atp6* (Fig. 7).

**Fig. 7.**
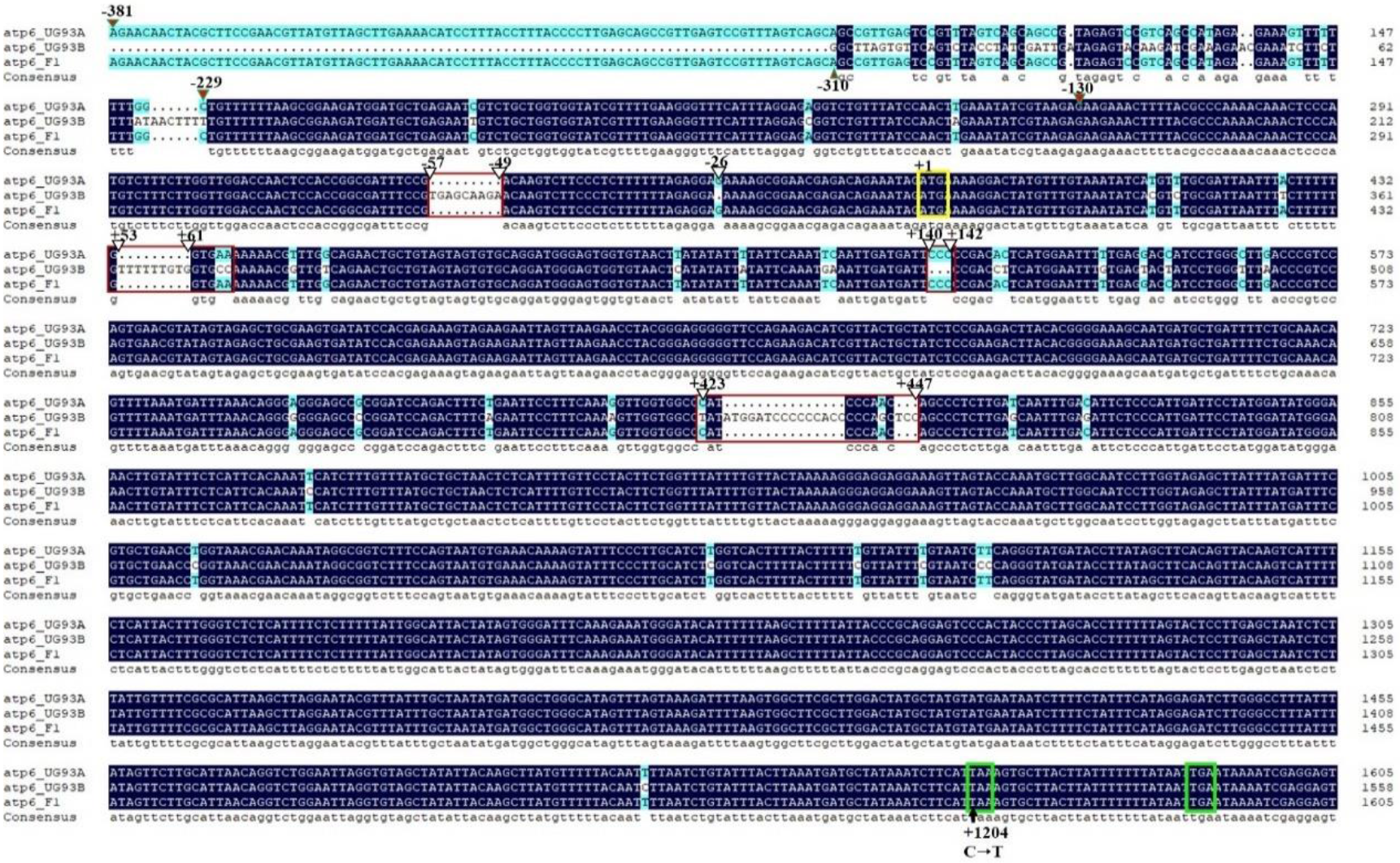
Transcription sequence analysis of *atp6*. The initiation codon and stop codon were indicated by yellow and green box, respectively. The variable regions among UG93A, UG93B and F_1_ were marked with red boxes, and the variable sites were indicated by open triangles. The transcripton initiation site were indicated by solid red triangles. The C-T editing site of the cDNA sequence was indicated by black arrow.

The 5’ - ends of *atp9* was identity in UG93A, UG93B and F_1_ (UG93A/UG93R), located at position −71 upstream of the start codon, but was different at 3’- end. The transcription termination site of the 3’ - end was located at +271 in UG93A, but +397 downstream of the stop codon in UG93B and F_1_ (UG93A/UG93R), which was consistent with the mRNA transcript among UG93A, UG93B and F_1_ (UG93A/UG93R) as previously research (Fig. S7). In the cDNA sequence of *atp9*, a C-T conversion was identified at the nucleotide position +262 of reading frame compared to *atp9* gDNA sequence, resulting in a generation of stop coding TGA, that lead to early termination of the CDS region of *atp9* (Fig. 8).

**Fig. 8.**
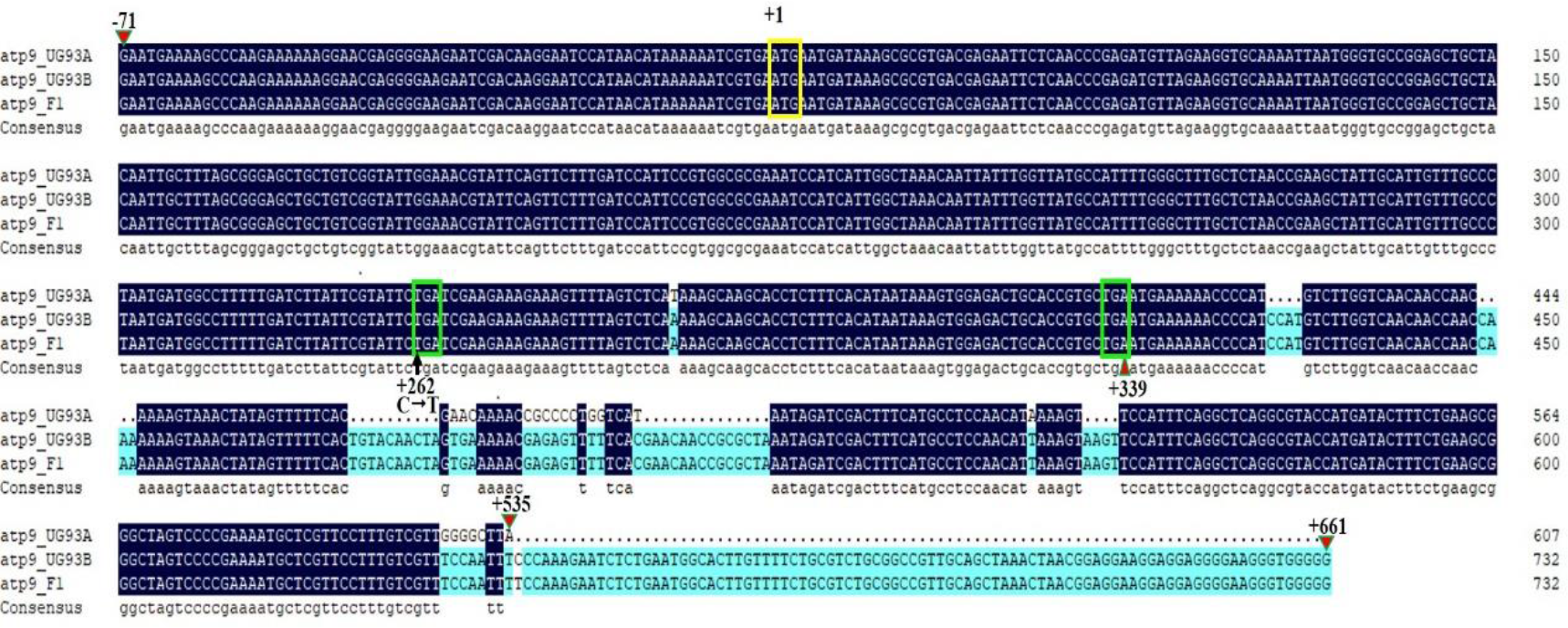
Transcription analysis of *atp9*. The initiation codon and stop codon were indicated by yellow and green box, respectively. The variable sites were indicated by solid red triangle.

The *cox3* RNA was shown to accumulate in two different forms with the 5’- ends located at −164 and −313 upstream of the start codon in UG93A. One 5’- ends mapped at the −311 in UG93B, and two 5’- ends located at nucleotide positions −313 and −442 upstream of the start codon. The 3’ - ends were mapped at the same position in the three cytoplasmic of kenaf (+314 relative to the stop codon) (Fig. S8, S9).

### The highly diverged of the mitochondrial promoter architecture in kenaf

The YRTA motifs are widely present in the mitochondrial promoters of monocot and dicot plant species. A search for analyzing the consensus sequences in kenaf mitochondrial promoter regions found that only eleven of them contained the YRTA motifs (Table 2.). Six of these core element sequences are located at the position of −4 to −7 upstream of the transcriptional start, and the other five are located at the position of −25 to −29 upstream of the transcriptional start. These promotor sequences were further analyzed with WebLogo software, which is designed to generate sequence logos within a multi-sequence alignment for sequence conservation at particular positions. Our result showed that the promoter sequences were less conserved overall, but the nucleotide of T or A were observed with highly frequency at the upstream of initiation codon (Fig. 9). Our data suggest that the promoter sequences in kenaf mitochondrial genome are highly diverged, and the YRTA motif is not an essential element for the kenaf mitochondrial promoter activity.

**Fig. 9.**
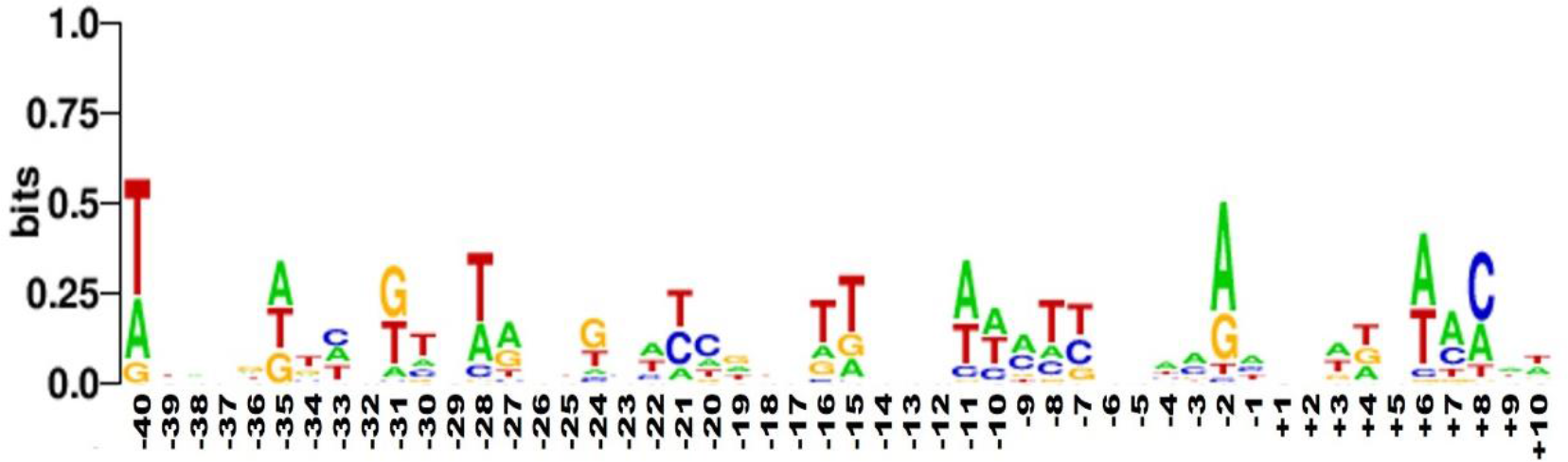
Summary of nucleotide sequences around the defined transcription initiation sites in kenaf mitochondrial genes (displayed in Table 2). Position +1 corresponds to the transcription initiation sites. The overall height of each stack indicates the sequence conservation at that position (measured in bits), whereas the height of each nucleotide within the stack reflects its relative frequency among the sequences.

**Table 2.**
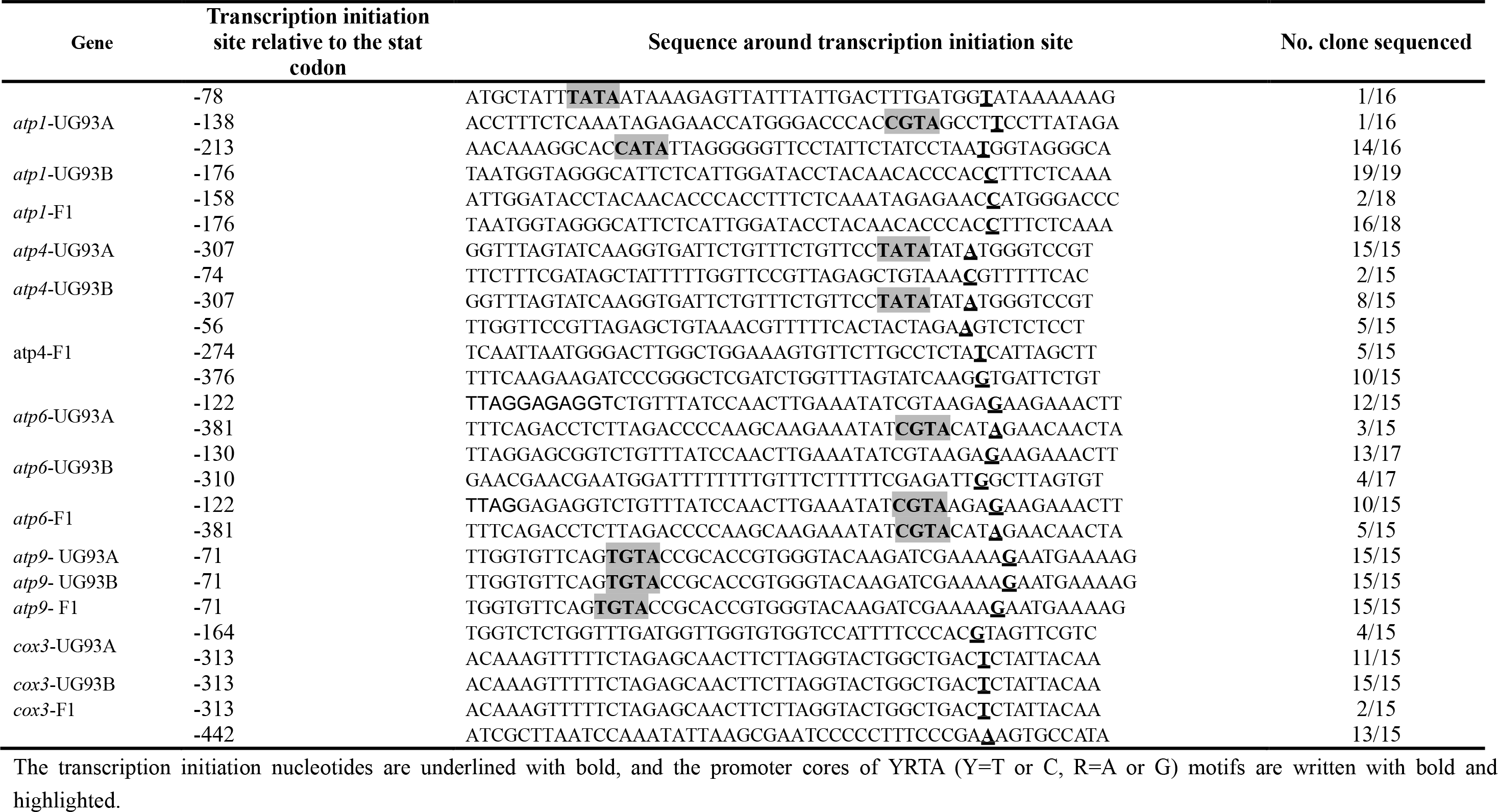
The transcription initiation sites and flanking sequences of the mitochondrial transcripts in kenaf.

## Discussion

### Comparative analysis of UG93A and UG93B mtDNAs

The evolution of the plant mitochondrial genome primarily includes genomic repeats, rearrangement, insertions, deletions, and nucleotide mutations among genomes (Clifton *et al.*, 2004; Tian *et al.*, 2006; Darracq *et al.*, 2010). In present study, a total of 398 SNPs and 230 InDels in UG93A compared to those in the reference mitochondrial genomic UG93B, far lower than those in *Brassica napus* and rice, which conserved 36,458 and 76,370 variations, respectively (Huang, 2013; Wang *et al.*, 2013). Since both CMS line UG93A and maintainer line UG93B were bred from wild-type UG93, belonging to cytoplasmic near-isogenic line. This line can be effectively eliminated the interference of redundant genetic information irrelevant to kenaf CMS. Hence these are ideal materials for understanding the molecular mechanism of kenaf cytoplasmic male sterility.

Plant mitochondrial genome is labile due to its recombination frequently among mitochondrial, chloroplast and nuclear DNA (Mower *et al.*, 2012). But most of the recombination sequences are located in intergenic regions and show a faster rate of evolution (Allen *et al.*, 2007). Furthermore, although many gene sequences were highly conserved between UG93A and UG93B, there were exceptions. The sequences of *rpl10*, *sdh4, cox1*, *rps14*, *cox2*, *atp4*, *rps7* between UG93A and UG93B were different, especially *atp6* that conserve active recombination region between UG93A and UG93B, which could be a good candidate genes for contributing to kenaf CMS phenotype. Finally, the number of SNPs between UG93A and UG93B mtDNA was aslo significant compared to those in a CMS line of sugar beet, which has 24 SNPs in 11 protein-coding genes compared to the fertile line (Satoh *et al.*, 2004).

SNPs and InDels are important factors that are responsible for biodiversity and are widely found in genome of plants and animals (Feltus *et al.*, 2004; Shen *et al.*, 2004; Mills *et al.*, 2006). In present study, most SNP and InDel variants occurred in non-coding regions, intergenic regions and introns of UG93A mtDNA and the distribution of InDels were mainly from 1 bp to 5 bp as identified in rice (Wang *et al.*, 2013) and *Chinese cabbage* (Park *et al.*, 2010). As InDels in coding region or promoter region are harmful to human health thus would be eliminated, while InDel variations between introns and intergenes need to be retained (Mills *et al.*, 2011). This suggested that variations in SNPs and InDels of genome could act a self-protection mechanism in process of biological evolution.

### Expression of mitochondrial genes in kenaf CMS

It has been documented that an altered transcription profile in CMS lines comparing to their maintainer lines and fertility-restored plants (hybrids) is one of the main features observed for CMS-associated genes (Hanson and Bentolila, 2004; Luo *et al.*, 2013). Many CMS genes have been proved to contain sequences similar to mitochondrial respiratory chain complex genes and co-transcribed with these genes, such as *atp1, atp6, atp8, atp9* and *cox2* (Hanson and Bentolila, 2004). In this study, we examined the transcript profile among CMS line UG93A, maintainer line UG93B and hybrid F_1_ by using of the 36 protein coding mitochondrial genes as RNA blot probes. A specific mRNA with *atp9* as probe was detected in UG93A, which has a smaller transcript size as compare to UG93B and F_1_ and distinct from other plants that CMS genes co-transcribed with adjacent essential genes to produce larger, dicistronic mRNAs (Liu *et al.*, 2007; Wang *et al.*, 2013). This data suggesting that it could be a novel regulation model to kenaf CMS, sequencing and functional analysis of *atp9* region is underway.

In present study, expression of *atp1* in UG93A and F_1_ (UG93A/UG93R) was lower compared with UG93B in the dicaryotic phase of anther development. Unlike expression of *atp1* in isonuclear alloplasmic CMS line P3A, P3B and F_1_ (P3A/992), which was lower than P3B and F_1_ (P3A/992). These current findings indicated that expression of *atp1* were different in various cytoplasms, probably due to inconsistent nuclear background. The expression of *atp6* was lower in CMS line and F_1_ (UG93A/UG93R) compared with UG93B and consistent with expression of *atp6* in P3A, which was downward than in P3B and F_1_ (P3A/992) (Zhao *et al.*, 2016). This showed that *atp6* may be involved post-transcription in the kenaf CMS line. The expression of *atp9* in UG93A was significantly lower than in UG93B and F_1_ (UG93A/UG93R), revealing its significance in development of microspores of kenaf. While the expression of *atp9* in *ramee* male sterile line was significantly higher than fertile line (Duan *et al.*, 2008), indicating that the regulation of *atp9* was different among various crops. However, the expression of *atp4* in UG93B was higher, which was contrary to transcript that had weaker signal comparing with UG93A, suggesting that the expression of *atp4* was regulated by post-transcriptional processing. In brief, the expression of kenaf mitochondrial genes was significant difference in various cytoplasms, suggesting that pollen abortion in kenaf was resulted from differential expression of subunit gene in mitochondrial respiratory chain complex, leading in a insufficient energy supplying to pollen development.

### Heterogeneity of the 5’- end in plant mitochondrial genes

Multiple promoters are common feature of mitochondrial genes in plants (Kuhn *et al.*, 2005). A total of 30 transcription initiation sites were found in 12 mitochondrial genes in *Arabidopsis*, while only one termination site located at the 3’ - end (Forner *et al.*, 2007). There were also multiple transcription initiation sites located upstream of *atp1*, *atp6* and *atp8* while only one termination site located at the 3’ - end in rice (Kazama *et al.*, 2013; Wang *et al.*, 2013). In this study, we found only one transcription termination site located downstream of *atp1, atp4, atp6* and *cox3* in various cytoplasms of kenaf, but multiple transcription initiation sites located at the 5’ – end, resulting in different lengths of 5’ UTR. This may be associated with translation of mitochondrial genes and regulated by nuclear genes (Koizuka *et al.*, 2003; Kazama *et al.*, 2008), as the RNA processing factor RPF2 having triangular pentapeptide repeat required for the 5’ - mRNA formation of higher plant mitochondria (Jonietz *et al.*, 2010). The initiation site at 5’- end of *atp9* was identical in UG93A, UG93B and F_1_ (UG93A/ UG93R), but 3’ - end was different, having 126-bp shortage in UG93A compared with UG93B and F_1_ (UG93A/ UG93R). The present results was consistent with the 5’ -transcription initiation site of *rps3*, which was same in both sterile cytoplasm and fertile cytoplasm of sugar beet having 460-bp shortage at the 3’ - transcription termination end of sterile cytoplasm compared to fertile cytoplasm (Matsunaga *et al.*, 2011).

Our analysis on the promoter architecture showed that most of the kenaf mitochondrial promoters were not conserved with the YRTA motifs. This is fairly differ from that in other dicot plants, which showed immediately upstream a CRTA core sequence (Kuhn *et al.*, 2005). The results showed that that the promoter sequences in kenaf mitochondrial genome are more diverged than those in other plant species, indicating that the YRTA motif is not an essential element for the kenaf mitochondrial promoter activity. However, A or T-rich elements immediately upstream of the promoters were present, which is similar to those observed in other plant mitochondrial gene promoters that have been shown to be essential for the function of the promoters (Zhang and Liu, 2006). Our data highlighted substantial diversity in transcriptional regulation of mitochondrial genes, and the molecular mechanism in regulation of mitochondrial gene transcription need further research for better understanding and confirmation in light of current findings.

## Conclusion

Understanding the molecular mechanism of CMS in kenaf is important for heterosis. Plant CMS is closely related to rearrangement and transcriptional regulation of its mitochondrial genome. Comparative genome and Northern blot shows that the CDS of *atp6* had active region of mitochondrial genome recombination and the transcript of *atp9* in F_1_ was regulated by nuclear genes, despite having the same cytoplasm with UG93A. The qRT-PCR showed that expression of kenaf mitochondrial genes was significant difference in various cytoplasms, suggesting that pollen abortion was resulted from differential expression of subunit gene in mitochondrial respiratory chain complex, leading in insufficient energy supplying to pollen development. CR-RT-PCR showed that there were multiple transcription initiation sites at the 5’- end of mitochondrial genes, revealing common feature of the 5’- end heterogeneity of mitochondrial genes in kenaf. Our result has provided the basic information for elucidating the mechanism of CMS in kenaf and hence can be used as theoretical reference for better understanding the transcriptional regulation of mitochondrial genes.

## Acknowledgments

This study was supported by the National Science Foundation of China (No. 31571719 and No. 31660430), the Chinese Postdoctoral Science Foundation (No. 2016M592608) and Natural Science Foundation of Guangxi Province (No. 2018JJB130045). The authors thank the Wuhan NextOmics Biotechnology Limited Company (China) for their help in resequencing the UG93A mtDNA of kenaf.

## Author Contributions

R. Z. initiated the experiment. X. L. conducted the experiment and drafted the manuscript. X. K. and B. Z. isolated the kenaf mtDNA. M. L. conducted the Northern blot analysis. M. W. and S. P. assisted with the experiment. A. K. and F. M. revised the manuscript and Y. Z. provided suggestions and edits for the manuscript.

